# The adaptability of the ion binding site by the Ag(I)/Cu(I) periplasmic chaperone SilF

**DOI:** 10.1101/2023.03.31.534560

**Authors:** Ryan M. Lithgo, Marko Hanževački, Gemma Harris, Jos J. A. G. Kamps, Ellie Holden, Justin LP Benesch, Christof M. Jäger, Anna K. Croft, Jon L. Hobman, Allen M. Orville, Andrew Quigley, Stephen B. Carr, David J. Scott

## Abstract

The periplasmic chaperone SilF has been identified as part of an Ag(I) detoxification system in Gram negative bacteria. Sil proteins also bind Cu(I), but with reported weaker affinity, therefore leading to the designation of a specific detoxification system for Ag(I). Using isothermal titration calorimetry we show that binding of both ions is not only tighter than previously thought, but of very similar affinities. We investigated the structural origins of ion binding using molecular dynamics and QM/MM simulations underpinned by structural and biophysical experiments. The results of this analysis showed that the binding site adapts to accommodate either ion, with key interactions with the solvent in the case of Cu(I). The implications of this are that Gram negative bacteria do not appear to have evolved a specific Ag(I) efflux system but take advantage of the existing Cu(I) detoxification system. Therefore, there are consequences for how we define a particular metal resistance mechanism and understand its evolution in the environment.

## INTRODUCTION

Silver compounds are effective antimicrobials that are highly toxic to many Gram negative bacteria including *Escherichia coli*. Silver is non-toxic to humans and other higher eukaryotes, except if ingested in very large quantities[1, 2] unlike other bactericidal metal ions such as mercury. As such, silver compounds can be found within the linings of bandages and as additives in creams, both of which are used in hospital burn wards, and as linings for medical equipment such as catheters[2, 3]. Silver compounds are have been used as an antimicrobial in a wide variety of household and personal products such as washing machine interiors, deodorants and some items of clothing[4–7].

With such a wide-spread use of an unlicensed antimicrobial and its unavoidable release into the environment, there has been an inevitable emergence of silver resistant bacteria. The first cases, reported in a US hospital burns ward in the 1960s, were of resistant *Salmonella enterica*[8], but silver resistance is being reported across a wide range of gram-negative bacteria. The archetype silver resistance genes reside on a 383 kb plasmid, pMG101.[9, 10] Studies of *E. coli* containing the plasmid pMG101 showed that the bacteria were able to survive and grow in the presence of 6x the normal lethal dosage of Ag(I)[10, 11]. Monovalent silver ions, Ag(I) are the active element, rather than metallic silver itself[12–14]. Therefore, the structural requirements for recognition of Ag(I) versus Cu(I) are of great interest to understand not only silver metal ion resistance, but also how proteins discriminate between these apparently very similar ions *in vivo*. There is a cluster of nine silver resistance genes (*sil*), *silABCEFGPRS* found in many Gram negative bacteria[1, 10, 11, 15, 16]. Gene products include SilABC, an RND+ efflux pump and membrane complex that spans the inner and outer membranes; SilP is an inner membrane Type P_1B_-ATPase; SilR and SilS form a two component signalling system that controls inducible silver resistance. The proteins SilE and SilF are periplasmic chaperones, while the role of SilG is so far unknown. Previously, we have characterised Ag(I) binding to SilE, showing that it is a disordered protein that folds upon binding 6 Ag(I), but can bind up to 8 ions[17]. We now turn our attention to the chaperone SilF

It is known from work on the copper resistance mechanism that the SilF homolog the periplasmic chaperone CusF binds Cu(I) and Ag(I), but not Cu(II)[18]. It is responsible for shuttling Cu(I) to the CusABC efflux complex for export out of the cell. Recent evidence has also emerged that CusF is prevented from oxidation of its ion binding methionine residues by MsrPQ, enabling it to remain active in the more oxidising environment of the periplasm[19]. It is likely that SilF performs a similar role as a metallochaperone[11].

In this paper we characterise structurally, biophysically and by simulation the relative Ag(I) and Cu(I) binding properties of SilF from *Escherichia coli* and show how measurement of the metal ion specificity illuminates the biological role of the *sil* system.

## Results

### The Structures of Apo and Holo SilF

The first 37 residues of SilF contain a periplasmic export sequence as well as a short predicted disordered region. Therefore, we cloned and expressed SilF from residues 38-120. The molecular oligomerization state of SilF_38-120_ was first investigated by size exclusion chromatography coupled to multi-angle light scattering (SEC-MALS) and analytical ultracentrifugation (AUC). Single species were observed by both techniques (Figure. S1). The molecular weight of SilF_38-120_ was determined to be 8.74 kDa (±7.9%) by SEC-MALS (Table S1) and 9.0 (± 0.2) kDa by sedimentation velocity analytical ultracentrifugation (SV-AUC). Both of these values are consistent with the calculated molecular weight of 9.1 kDa, indicating that SilF_38-120_ is monomeric in solution; neither method detected any higher order aggregates. Next, we determined the structure of SilF_38-120_ using X-ray protein crystallography (Figure 1A). Apo-SilF38-120 packed in a hexagonal unit cell with one protein chain per asymmetric unit. The protein has a β-barrel topology composed of five β-stands arranged in the β1-β2-β3-β5-β4-β1, similar to that observed for Cu(I) chaperone CusF[20] (Figure 1D). Unlike CusF, SilF_38-120_ has an additional 15 amino acid α-helix positioned between strands β3-β5. In CusF this is an extended loop with no discernable helical or sheet secondary structural elements (See Figure 1D). Interestingly, the prediction from AlphaFold2 (Figure 1E), which will have been trained using CusF, but not our structures, does show a shorter helix around 50% of the size of the one observed in SilF_38-120_ in this position; the rest of the residues are predicted to be disordered.

**Figure 1.**
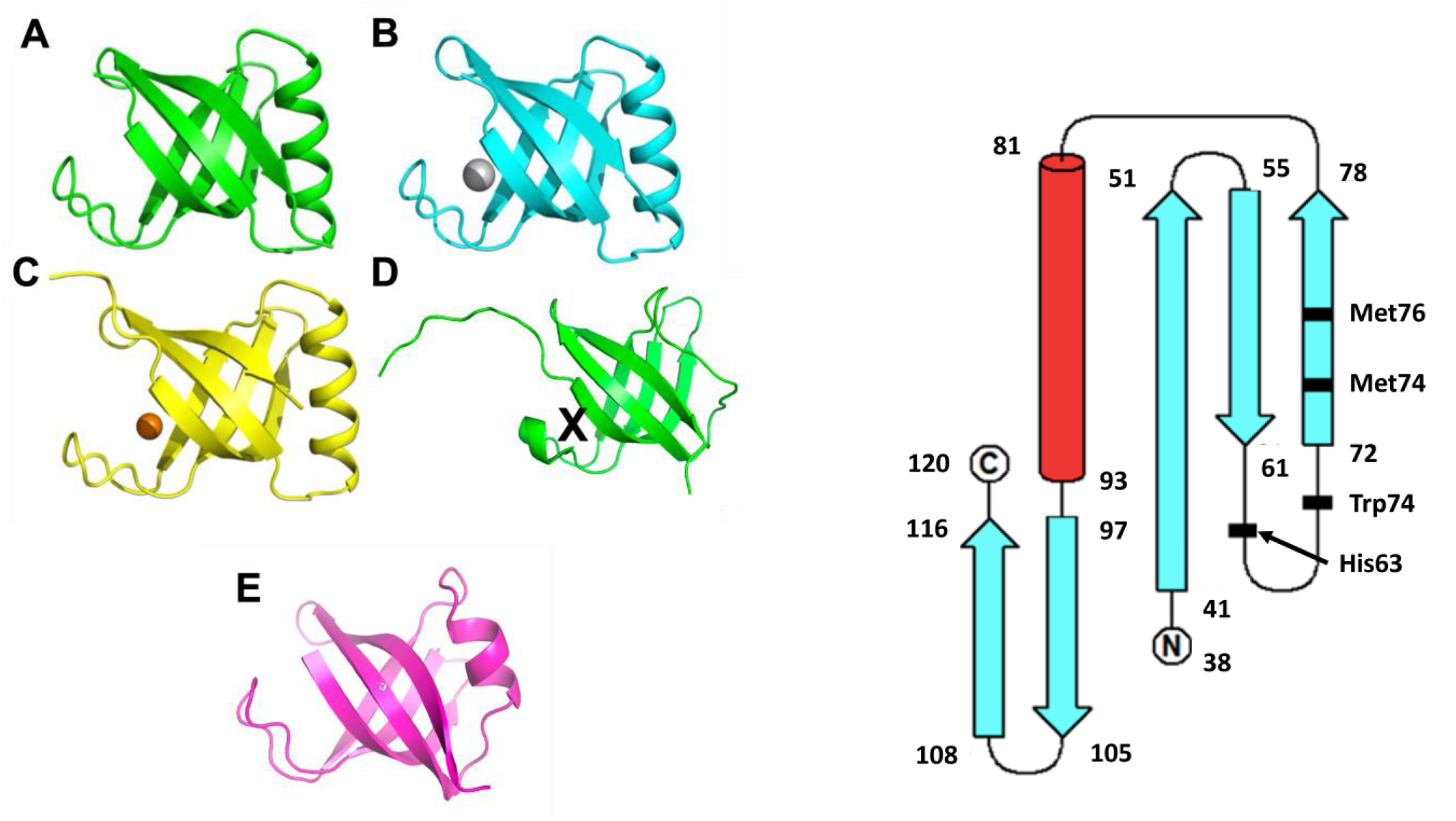
Ribbon diagram of **A:** apo- SilF_38-120_ (PDB: 8BBZ) **B:** Ag(I)- SilF_38-120_ (PDB: 8BHU) **C:** Cu(I)- SilF_38-120_ (PDB: 8BI1) and **D:** CusF (PDB: 2VB3). The ion binding site in CusF is shown by X. **E:** AlphaFold2 prediction of SilF. The prediction for the structure shown is at the level of pLDDT > 90 %; the predicted disordered signal peptide has been omitted. **F:** Topology of SilF showing the residues involved in ion binding marked in bold.

Co-crystallization of SilF_38-120_ with either Ag(I) or Cu(I) under anaerobic conditions yields an orthorhombic unit cell with three protein chains per asymmetric unit (Table S2) and each chain with one metal ion bound (Figure 1B and 1C) close to one end of the β-barrel (Figure 1 and 2). We attempted co-crystallization with Cu(II) but were unsuccessful. Calculation of the sedimentation coefficient using the coordinates of each of the monomers[21–23] taken from the respective crystal structures yielded a value of 1.25 *S*, consistent with a monomer of SilF_38-120_ measured by sedimentation velocity. In both structures the metal ion is tetrahedrally coordinated (Figure 2), and metal ion binding appears to have a limited effect on the overall conformation of the protein. Root mean square deviations (RMSD’s) derived from the C_α_ atoms are 0.99 Å for Ag(I)/Apo and 1.09 Å for Cu(I)/Apo. Larger deviations are observed primarily in the metal binding site, the top of the α-helix and the final loop that leads into the C-terminus. Re-calculating RMSD’s with these two regions missing reduces the value to 0.68 Å for Ag(I)/Apo and 0.98 Å for Cu(I)/Apo, showing that changes in flexibility is confined to the binding site and these regions. There are distinct conformational differences of residues at the metal ion binding site, and the loop connecting β4-β5. In both the Ag(I) and Cu(I) bound, structures the metal ion is coordinated with distorted tetrahedral geometry through the donor groups NE2 of His63 and the two thiol groups of Met74 and Met76. Figure 2 (both panels and Table S3) show the bonds and their lengths between the residues and metal ions for both SilF_38-120_. A comparison with CusF[18] (PDB: 2VB3, Table S3) shows that the metal ion coordination distances are identical. In SilF_38-120_, Trp71 acts as a cap over the metal coordination site, which likely further stabilises binding of the Ag(I) via cation-π interactions between the aromatic indole ring system and the positively charged metal ion.

**Figure 2:**
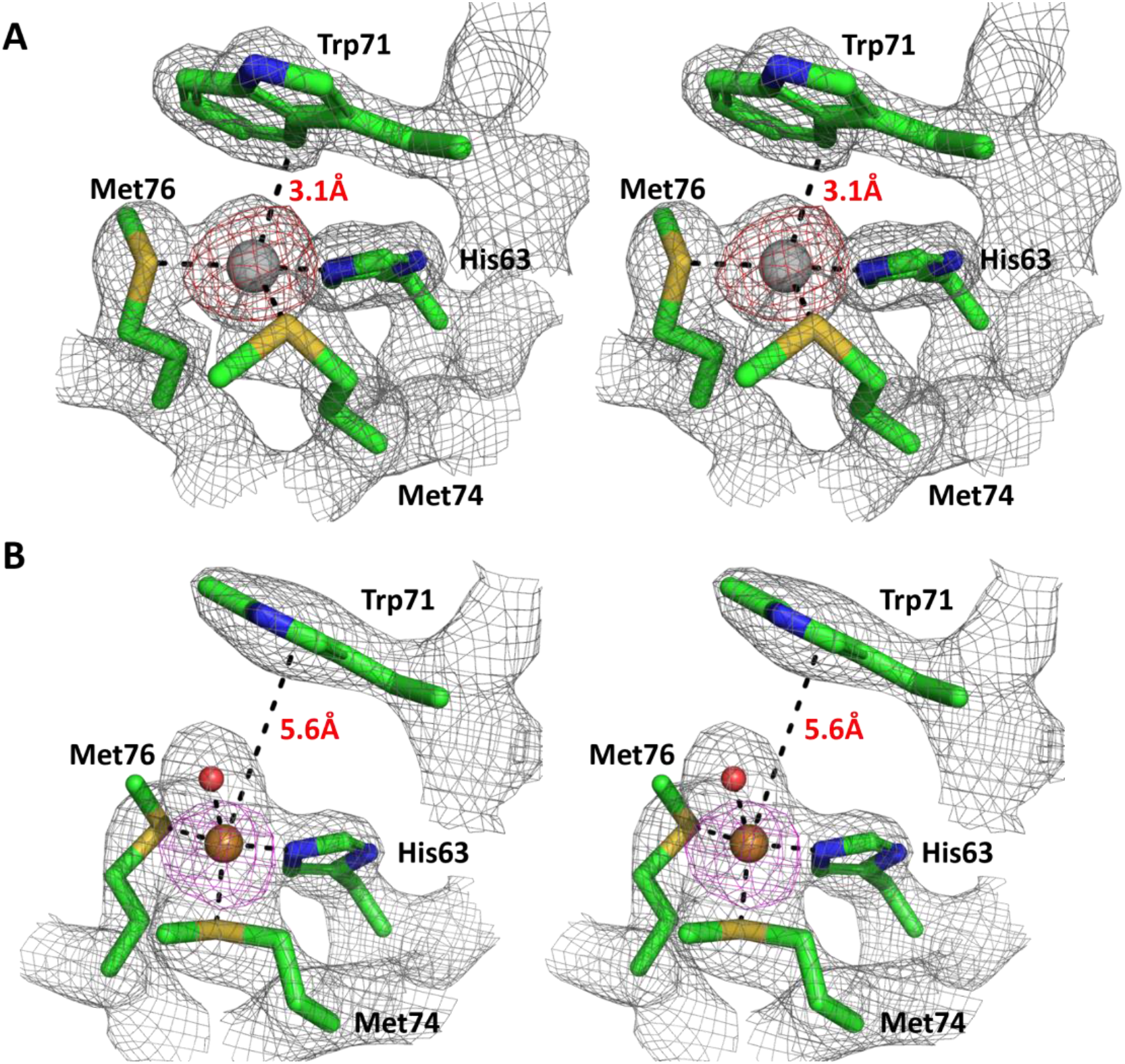
Wall-eye stereo view of cation binding site in SilF_38-120_. Residues involved in **A:** Ag(I) and **B:** Cu(I) binding in SilF_38-120_. Electron density at 1.5 σ from refined 2F_0_-F_c_ maps is overlaid for information; red is the anomalous density. The Cu(I) ion is smaller (0.60-0.74 Å) than the Ag(I) (1.0-1.14 Å) and as such an extra water molecule is coordinated in the binding site.

The Cu(I) coordination geometry within the binding site is noticeably different that of to Ag(I) where the fourth coordination position of the Cu(I) ion is occupied by a water molecule. The Trp71 is now blocked from directly interacting with the metal. Furthermore the side-chain of Met74 adopts a different rotameric conformer compared to the apo and Ag(I) structures. Such flexibility allows the protein to provide ligands at shorter coordination distances so as to accommodate the smaller ion. By contrast, the conformations of Met76 and His63 remain unchanged between structures. This difference in the observed water coordination for SilF_38-120_-Cu(I) binding is again notably different to CusF-Cu(I) binding which demonstrates close coordination of the equivalent Trp. It is argued that this interaction and connected water exclusion from the binding side is important to prevent Cu(I) oxidation when bound to CusF[20, 24–26].

Structural homology searches[27] revealed that, in addition to CusF, the subunit S1 of Pertussis toxin (PDB 1PRT), a domain from pro-protein glutaminase (PDB 3A54), and subunit B of subtilase cytotoxin (PDB 3DWA) were most similar in structure to SilF_38-120_. These structures are either domains or small proteins classified as oligonucleotide/ oligosaccharide binding (OB) fold proteins that are typically found in oligonucleotide/ oligosaccharide binding domains. They are comprised of 5 or more β-strands interlinked with either an α-helix, extended loop or a three-helix bundle between strands β3-β4[28–30]. The binding regions of OB-fold proteins vary with no singular defined binding region with different proteins using different loops at either end of the barrel to bind their target ligand[31, 32]. The molybdenum sensor ModE that regulates transcription of several genes involved in cellular molybdenum homeostasis, is currently the only other example of a metal binding OB-fold protein[33], although here metal binding is a prerequisite for binding ssDNA[34]. SilF and CusF are therefore the only examples of OB-fold proteins with the sole function of metal ion binding.

### Conformational Flexibility

Hydrogen-deuterium mass spectrometry (HDX-MS; Figure S2) was used to probe changes in conformational flexibility over time (30s, 5min & 30min) and to provide localised in-solution evidence of metal binding to SilF. A coverage map was generated, covering 88.9% of the protein amino acid sequence (Figure 3 and S2).

**Figure 3:**
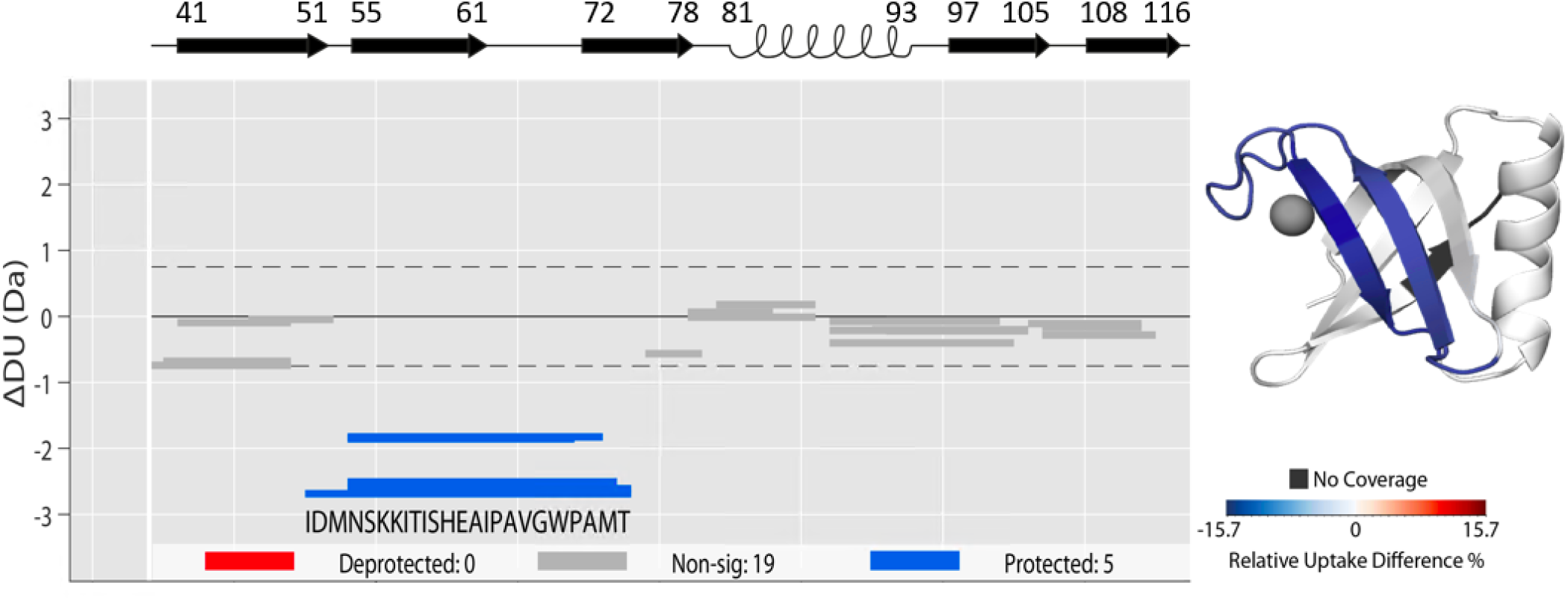
Changes in peptide deuterium uptake over time measured by HDX-MS for the Apo SilF_38-120_ versus Ag(I)- SilF_38-120_; experimental conditions are detailed in the supplementary information. **Left** Wood differential plot showing statistically relevant changes (hybrid significance testing p<0.001) in deuterium uptake after 30 mins of incubation in D_2_O. **Right** Wood differential overlaid onto SilF_38-120_ structure. Blue indicates a decrease in deuterium uptake, red for an increase. Changes are confined to the β sheet region and the binding site

Upon incubation with Ag(I) there are significant changes in deuterium uptake on peptides that span the amino acid sequence IDMNSKKITIS**H**EAIPAVG**W**PA**M**T (residues 52-75), which encompasses the metal ion binding residues His63, Met74, and pi-cation interaction with Trp71 (shown in bold. See also Figure 3 and S2). The observed reduction in relative deuterium uptake (15.7%) indicates that upon binding, Ag(I) interacts with amino acids within this region of the protein, in agreement with the crystal structure, and blocks the amino acids from exchanging with solvent deuterium. Due to the experimental setup we were unable to carry out Cu(I) binding in suitable anaerobic conditions.

To assess any changes in secondary structure that could occur in ion binding we employed synchrotron radiation circular dichroism spectroscopy (SR-CD).

There were distinct differences in SR-CD spectra in both the far and near UV regions (Figure S4A and B). Decomposition of the far UV region into secondary structure elements (Figure 4) showed that upon addition of both metal ions, there is an increase in α-helical content from 6 % to 16 % at the expense of disordered protein. Although changes in RMSD between the structures are observed (see above), there is little apparent change in helicity, although this may be due to crystal packing. Near-UV CD spectra, measuring the impact on aromatic residues, showed that upon addition of both Ag(I) and Cu(I) there were large spectral differences consistent with changes to Trp71 in the metal binding site (Figure S4B). It is noticeable that the Ag(I)/SilF_38-120_ complex has a near-UV CD spectra that is distinct from the Cu(I)/SilF_38-120_ complex. This we attribute to the differences in binding mode of the Trp71 residue seen between the two crystal structures.

**Figure 4:**
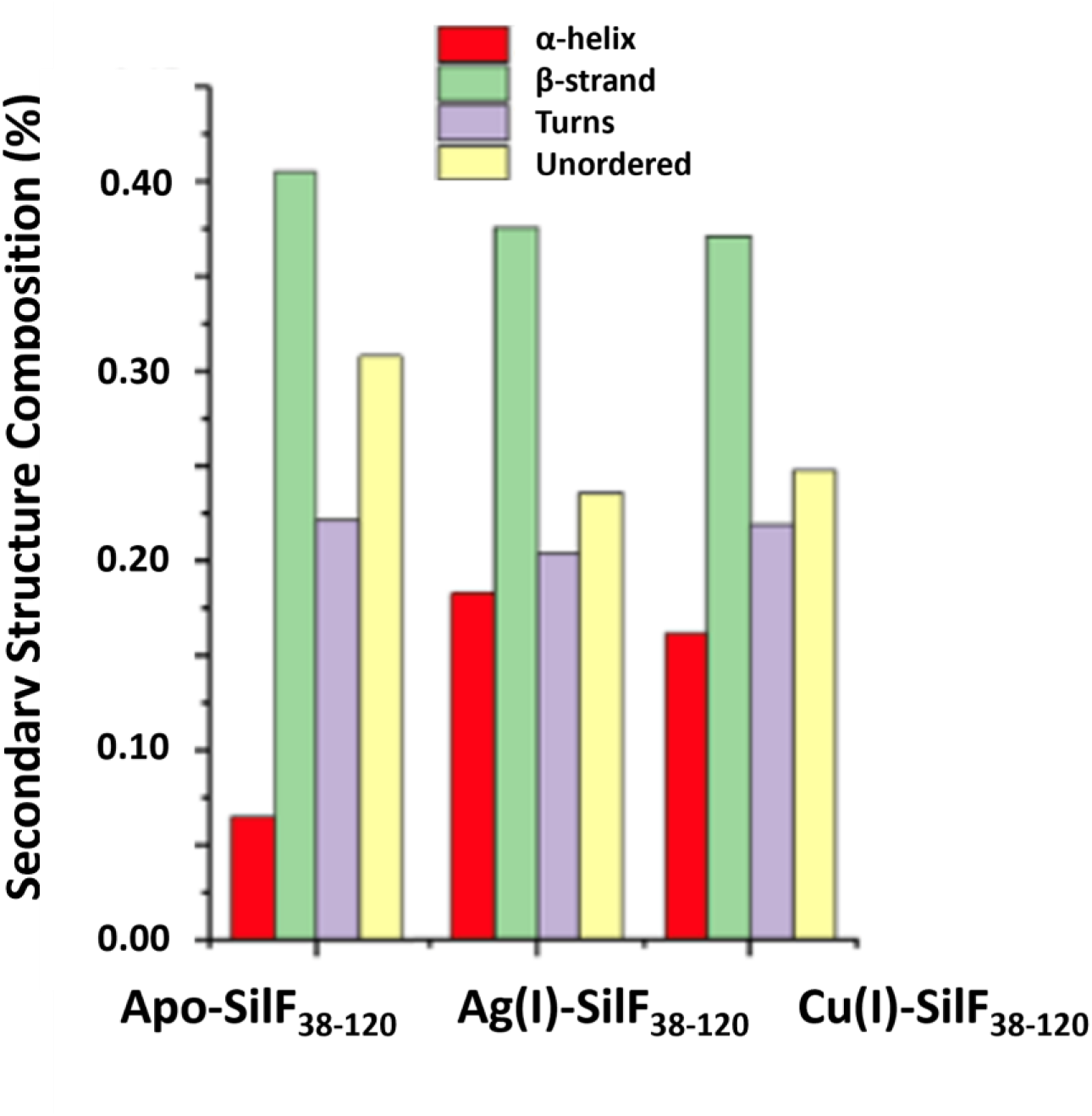
Results of secondary structure deconvolution of synchrotron radiation circular dichroism (SR-CD) of SilF_38-120_ in the apo, Cu(I) and Ag(I) bound forms. Experimental conditions are in the supplementary materials.

### Metal ion binding and specificity of SilF

In order to assess the affinity and thermodynamics of metal ion binding, we performed istothermal titration calorimetry (ITC) measurements. Studying binding events of Cu(I) in aqueous media is challenging for two reasons: i) Under aerobic conditions, Cu(I) readily oxidises by reacting with O_2_ from air, to give Cu(II); ii) under anaerobic conditions, Cu(I) undergoes a disproportionation reaction, resulting in the formation of Cu(0) and Cu(II) (equation 1)[35].

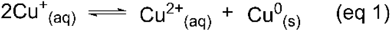

An excess of NaCl in solution can prevent the disproportionation of Cu(I) from occurring under anaerobic conditions, thus we performed all Cu(I) titrations at 1 M NaCl[36]. This equates with what was previously used for CusF/Cu(I) titration[37]. Solubility of AgCl is poor, resulting in precipitation even in presence of moderate concentrations of NaCl, preventing the use of identical salt concentrations for both experiments. Interactions of Cu(I)/Ag(I) with buffer molecules are considered to be weak. However, due to the relatively high concentration of buffer compared to the binding metal, a non-negligible effect of the buffer molecules on binding was observed, as has been reported previously for such ion titrations[38, 39]. To avoid these issues, the titrations were performed in absence of buffer.

We found that the binding of either metal to SilF_38-120_ has a 1:1 stoichiometry, confirming the observations from the crystal structures of a single binding site. Binding was exothermic with a small entropic penalty, which accords with the changes in flexibility and helicity seen in CD measurements. Ag(I) has a dissociation constant (*K*_d_) of 7.6 nM, whereas Cu(I) binds with a *K*_d_ of 30 nM: both determined free energies values were within experimental error of each other meaning that the affinity of SilF_38-120_ for either metal ion are very similar in size. The data is summarised in Table 1 and Figure S4 shows the thermograms from the ITC for each metal ion. Previous investigations using ITC of CusF under anaerobic conditions observed that Cu(I) was bound considerably more weakly than Ag(I)[37]. These experiments also yielded low estimates for stoichiometry of binding (0.52 for a protein known to bind with 1:1 stoichiometry), which indicates that non-specific buffer conditions were present in these measurements.

**Table 1:** Thermodynamic parameters derived from isothermal titration calorimetry experiments for SilF_38-120_ binding with Ag(I) and Cu(I). Data for CusF affinity and stoichiometry is taken from[37]: no other thermodynamic data was given in this reference.

| Expt | $K_d$<br>(nM) | Stoichiometry<br>($n$) | $\Delta H^\circ$<br>(kJ/mol) | $\Delta S^\circ$<br>(J/mol/K) | $\Delta G^\circ$(kJ/mol) |
| --- | --- | --- | --- | --- | --- |
| SilF<sub>38-120</sub> /Ag(I) | 7.6<br>($\pm 1.3$) | 0.99 ($\pm 0.03$) | - 55.1<br>($\pm 1.59$) | - 29.0<br>($\pm 1.7$) | - 46.4<br>($\pm 0.5$) |
| SilF<sub>38-120</sub> /Cu(I) | 30.0<br>($\pm 6.5$) | 0.96 ($\pm 0.22$) | - 54.8<br>($\pm 1.1$) | - 34.0<br>($\pm 3.9$) | - 44.6<br>($\pm 3.7$) |
| CusF/Ag(I) | 38.5<br>($\pm 6.0$) | 0.52 ($\pm 0.08$) | | | |
| CusF/Cu(I) | 495<br>($\pm 260$) | 0.82 ($\pm 0.09$) | | | |

### Molecular Dynamics and QM/MM analysis

To further probe the changes in conformation upon metal ion binding and computationally investigate differences in binding affinity of the different metal ions, classical molecular simulations and Quantum Mechanics / Molecular Mechanics (QM/MM) hybrid methods were employed. We used our determined crystal structures for Apo-SilF_38-120_, Ag(I)-SilF_38-120_ and Cu(I)-SilF_38-120_. We were unable to investigate a Cu(II)-SilF_38-120_ binding due to the previously described failure to produce crystals for this complex. Initial QM/MM optimisations matched well the structural differences observed for the different metal ion binding, including the addition of a water molecule in the coordination of Cu(I). Subsequent long (1.6 µs) timescale molecular-dynamics (MD) simulations confirmed conformational changes upon binding as seen in the HDX-MS results as demonstrated by principal component analysis (Figure 5A). There were changes in flexibility seen in the α-helix correlating well with changes in secondary structure contents indicated by SR-CD (Figure 4), again indicating that the lack of changes in helicity observed in the crystal structure may arise from crystal packing. Visualization of the dominant principal components and the helicity analysis from the simulations shown in Figure 5A-C show changes in conformation, as previously observed with HDX-MS and SR-CD. Analysis of the metal ion binding site from those simulations also confirmed the longer stability of the binding geometry and the differences between Ag(I) and Cu(I) binding. As shown in Figure 6 by the radial distribution Cu(I) retains tightly bound water molecules in its tetrahedral coordination sphere throughout the whole simulation. In comparison, whilst interactions of Ag(I) with water molecules could also be observed they appeared at a longer distance which allows Trp71 to bind closer to the metal.

**Figure 5A:**
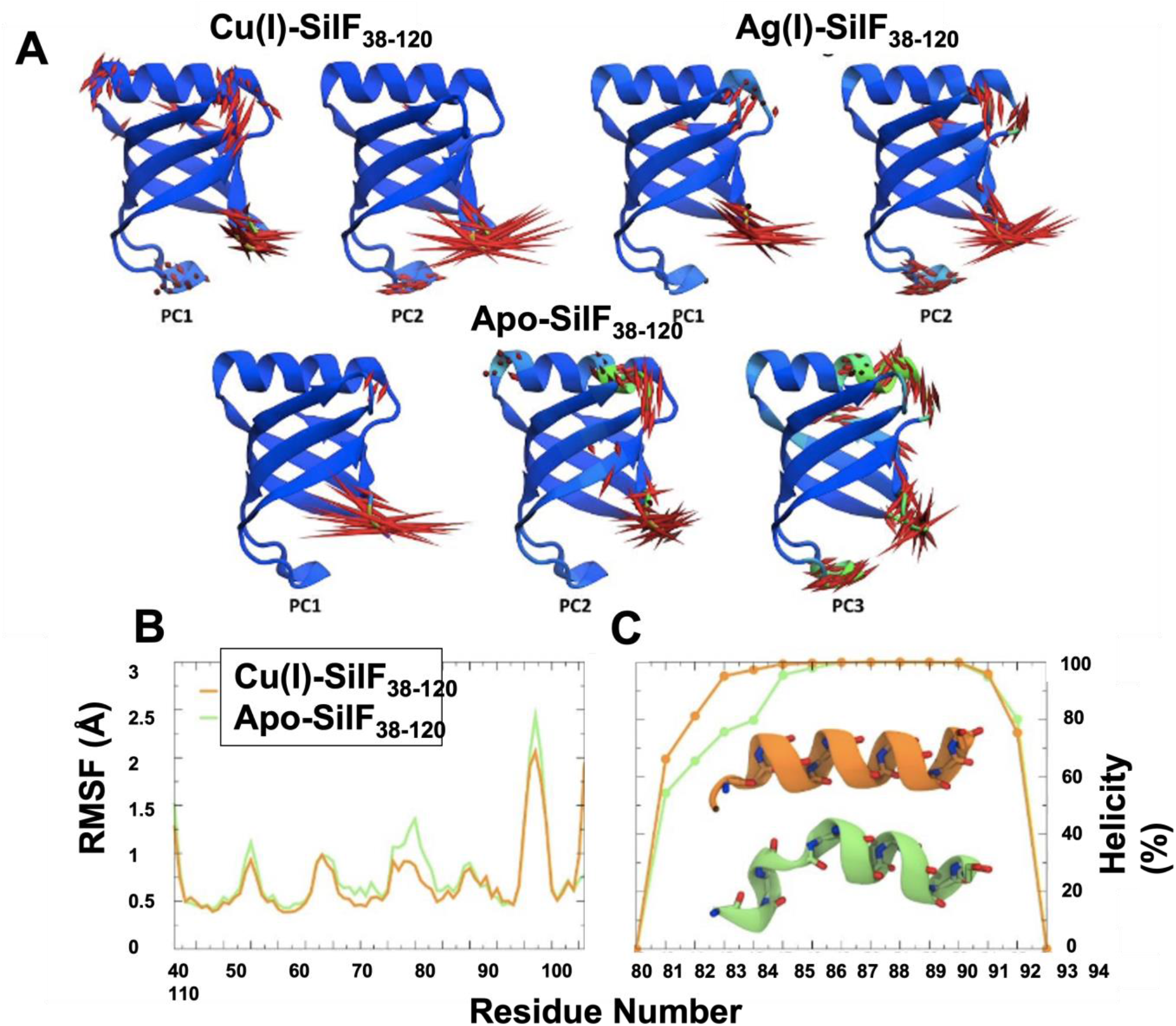
Normal mode displacements (larger than 2 Å) of first most prominent principal components. Displacements indicated in red by magnitude, those with high displacement along the helical axis in PC2 and PC3 of apo-SilF_38-120_ are indicated in green. The data was collected from 1.6 µs classical MD simulations of Cu(I)-SilF_38-120_, Ag(I)-SilF_38-120_and apo form. **B:** Residual root-mean-square fluctuation (RMSF) of SilF_38-120_ backbone atoms (N, Cα, C, O) in Cu(I)-SilF_38-120_ and apo-SilF_38-120_ form calculated from the reference X-Ray structure of apo protein. **C:** Secondary structure features of α-helix calculated for Cu(I)-SilF_38-120_ and apo form calculated using the database of secondary structure assignments (DSSP) algorithm, which assigns average secondary structure propensities over MD frames for each residue based on backbone amide (N-H) and carbonyl (C=O) atom positions. The protein was truncated in the simulations at the N and C termini to minimize fluctuations. All simulation data was used to generate the RMSF for each system, a total of 1.6 µs per system.

**Figure 6:**
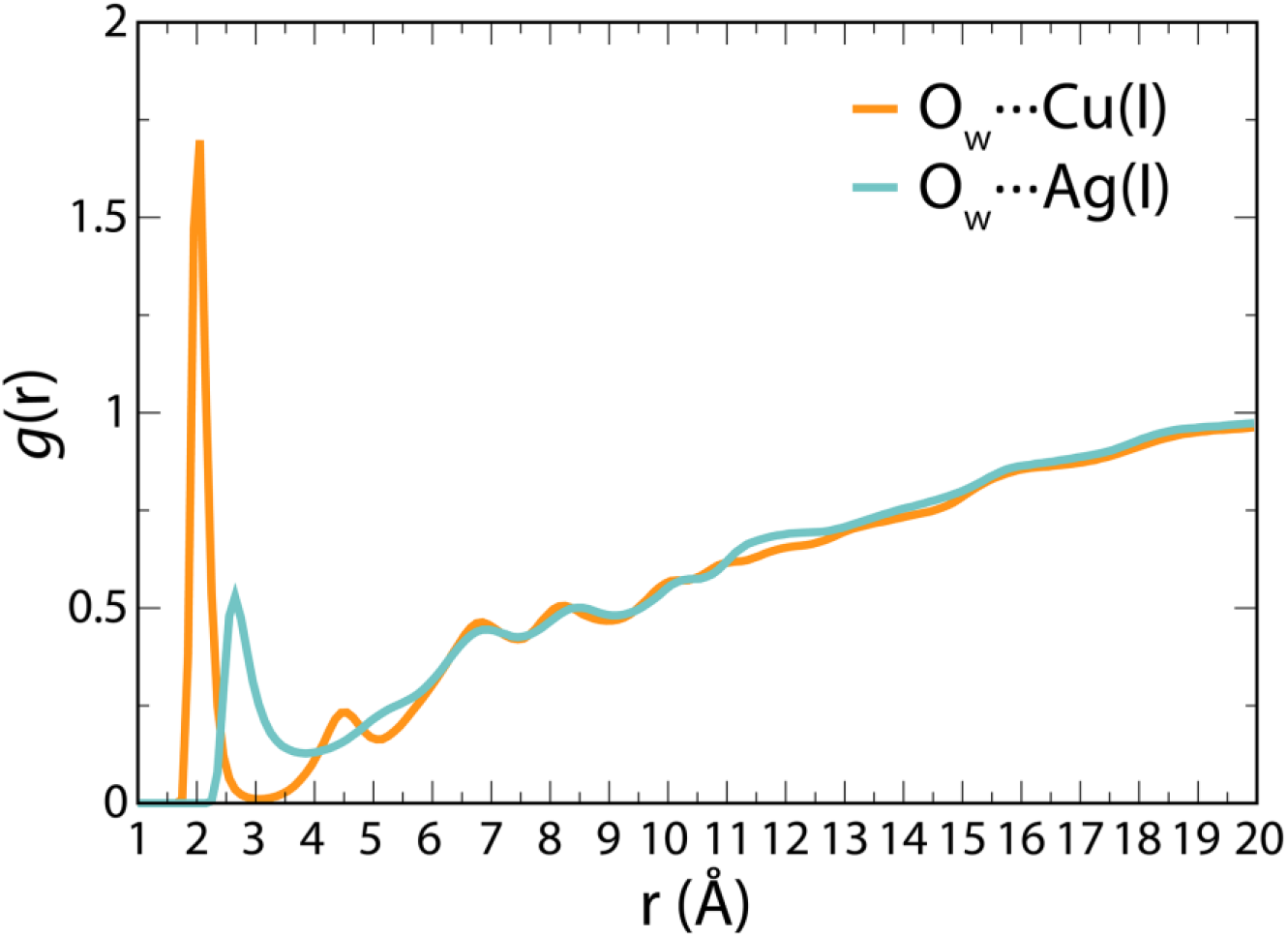
Radial distribution function (RDF) of water oxygen atoms around metal ions. The data was collected from 1.6 µs classical MD simulations

In contrast, the crystal structure of Ag(I)-CusF does not display a bound water molecule[20]. However when we carried out a molecular dynamics simulation of CusF-Cu(I) this showed a water molecule present coordinating between Trp71 and Cu(I) (see Figure S5) as seen in our Cu(I)-SilF_38-120_ crystal structure (see Figure 2). Previous simulation studies were able to replicate the absence of this water molecule in CusF by re-parametrising the Lennard-Jones potential to increase the strength of the Cu(I)-π interaction. When this study used a traditional potential, 60 % occupancy of water akin to what is seen in the Cu(I)-SilF_38-120_ structure was observed[40]. Coordination between Trp71 and Cu(I), as shown in Figure S5 and as demonstrated by the sharp peak in the radial distribution function of water around the metal ions, is presented in Figure 6.

The residency times for water in the first hydration shell around the binding site of Cu(I)-SilF_38-120_ is on average 101 ps with a standard deviation of 117 ps. However, for Ag(I)-SilF_38-120_ this value was considerably shorter with an average of 5 ps and a standard deviation of 8 ps. (Solvation shells are defined as the area under the first solvation peak as seen in the radial distribution function in Figure 6). Therefore, judging from the combination of those different sets of simulations and the crystallographic evidence of the two binding sites, it appears that the binding site of SilF is adaptable with at least two different possible modes of binding including the observed water coordination, differently to the CusF binding site where water exclusion dominates.

To further probe the changes in SilF protein conformation upon metal binding, multiscale modelling was applied based on structures from the extensive classical molecular dynamics (MD) simulations. For this analysis over 40 ps of QM/MM simulations starting from equilibrated MD simulations were performed.

Those simulations demonstrated stable ion binding coordination and ten snapshots have been picked to calculate the binding enthalpies of the different ions relative to their solvation enthalpies in water (see Supporting Information for methodology). The binding enthalpies calculated with static QM/MM methodology demonstrated large variations, and although a slightly higher average binding enthalpy for Ag(I) by 11.8 kJ/mol is calculated, (Figure 7), the standard deviation of each of the enthalpy calculations is around three times this value, making this enthalpy difference between the two binding events statistically insignificant. This is also in line with the experimental ITC measurements (see Table 1) where changes in enthalpy and free energy changes cannot be distinguished statistically for both ions.

**Figure 7:**
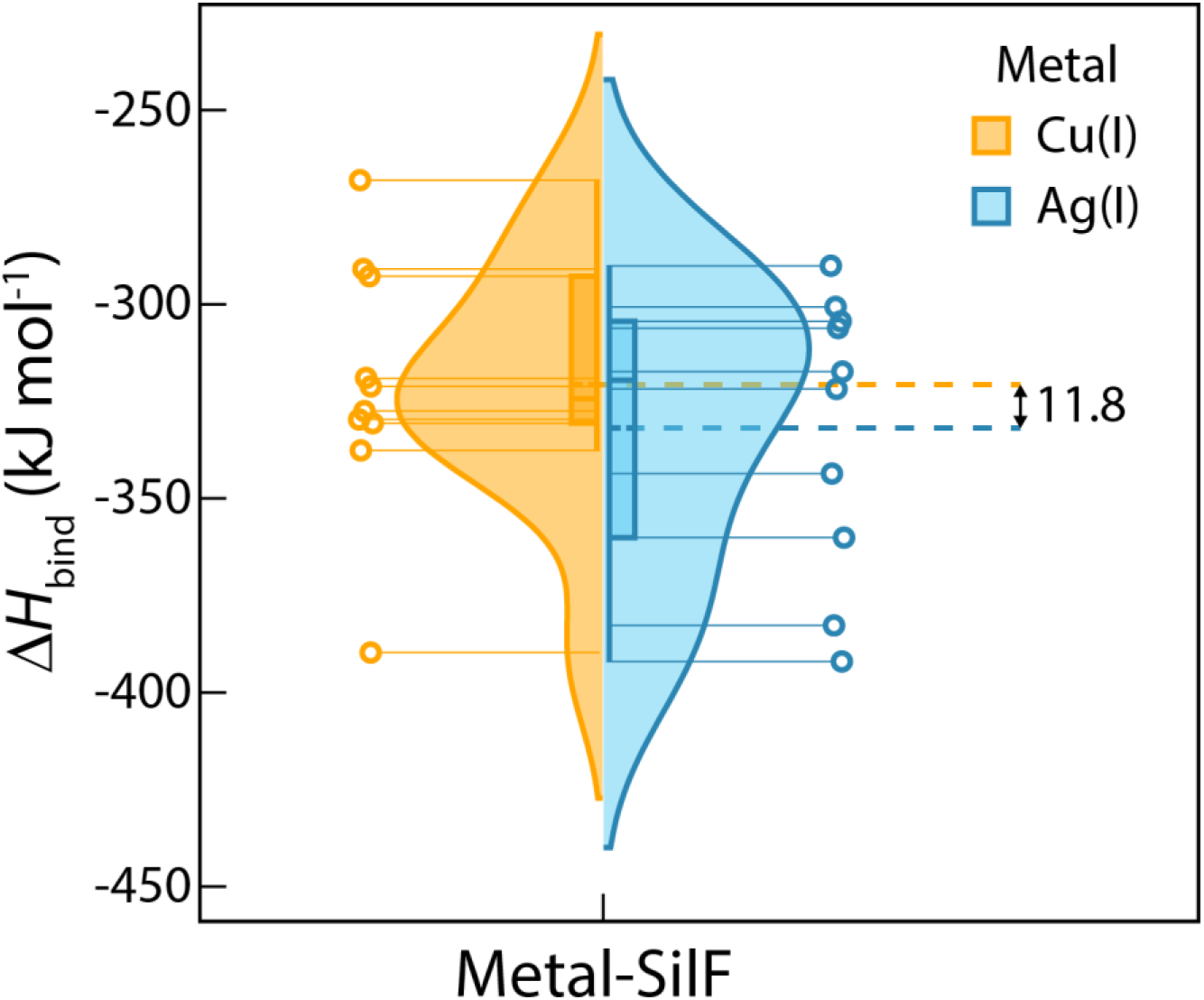
Violin plot representation with box and discrete data points of binding enthalpies for Cu(I)-SilF_38-120_ and Ag(I)-SilF_38-120_ obtained from ONIOM calculations using ten different structures from QM/MM MD simulations. Although a slightly higher enthalpy is calculated for Ag(I) binding, this is not significant within the wider spread of calculated enthalpies where the standard deviations for Cu(I) and Ag(I) binding were 32.86 kJ/mol and 35.88 kJ/mol, respectively. A violin plot contains box plot (median, interquartile range and upper/lower half shown as orange and blue lines inside the box for Cu(I) and Ag(I), respectively) with the addition of a rotated kernel density plot for each system. The quartiles Q1 and Q3 are computed using the linear interpolation method. A difference between average values of Δ*H* for Cu(I) and Ag(I) is depicted with dashed lines.

## DISCUSSION

Biophysical and biochemical analysis of proteins comprising the *sil* gene cluster has been limited to date, with sequence homology used to infer function, and the proposed resistance mechanism, after comparison with the more extensively studied *cus and cue* systems[1, 10]. Such analysis is consistent with the model that the SilF is a periplasmic metal-binding chaperone capable of binding both Ag(I) and Cu(I) ions and shuttling them to the SilCBA complex to aid metal ion detoxification[41, 42].

Our analysis supports that the observed lack of any appreciable metal ion binding preference of SilF for Ag(I) compared to Cu(I) which was evident from the ITC measurements. The dissociation constant of 7.5 nM for Ag(I) is a little tighter than the affinity measured for CusF (38 nM)[37], whereas the relative Cu(I) affinities of SilF (30.0 nM) and CusF (450 nM) differ by more than an order of magnitude[15, 37]. The significant difference is in the binding stoichiometries, 0.5 for CusF(see [37]), 1.0 for SilF (this study) despite the crystal structures of both showing a single ion binding to the monomer.

The X-ray crystal structures show that the metal ion binding site is formed from strictly conserved residues His63, Trp71, Met74 and Met76 located at one end of the β-barrel. Upon Ag(I) binding histidine and methionine residues occupy three of the coordination sites of the bound metal. The coordination sphere is completed by the indole ring of Trp71, which forms a π-cation interaction to complete the tetrahedral coordination sphere. On binding of Cu(I) His63, Met74 and Met76 again adopt a distorted trigonal coordination geometry, but the fourth coordination site is occupied by a water molecule, preventing the formation of the π-cation interaction. Molecular dynamics shows that this persistence of the water molecule suggests the π-cation interaction with the Cu(I) ion is insufficiently stable to displace the solvating water molecule. Simple thermodynamics arguments are not enough to explain the presence of the water molecule since it does not persist in the CusF-Cu(I) structure, which has a highly similar active site architecture.

The inability of SilF to form a π-cation interaction with Cu(I) will decrease the binding enthalpy for copper, a further reduction in binding enthalpy will occur from a preference of sulphurous ligands to bind Ag(I) relative to Cu(I)[43]. The high-resolution structure of CusF with Ag(I) bound[18, 20] shows the coordinating methionine residues can adopt multiple conformations while still interacting with the metal ion. Such freedom of movement within the relatively flexible coordination sphere of Ag(I) reduces the entropic penalty of metal binding. The smaller ionic radius of Cu(I) (0.60-0.74Å) relative to Ag(I) (1.0-1.14Å) constrains the geometry of the coordinating methionines as they tuck into the binding site and interact with the metal. Stabilisation of Cu(I) via a π-cation interaction has previously been demonstrated for Cu(I) binding to CusF[18], and is likely the cause of the difference in Cu(I) binding affinity between CusF and SilF. The adaptability of the binding site leading to this lack of ion specificity is also supported by results from QM/MM binding enthalpy calculations.

Displacement of the tryptophan loop is not the only conformational change observed within SilF. HDX-MS shows changes globally across the protein and CD spectroscopy show an increase in alpha helical content of SilF upon cation binding. Although we do not see appreciable changes in helix formation from the crystal structures, most likely due to crystal packing forces, both of the solution experimental observations are well supported by the simulations where changes in helicity are clearly over long simulation times between the apo and holo SilF. Together these show that the capping helix is unstable in the apo-protein and indeed could be unfolded to a degree in solution, as observed for CusF, with metal binding stabilising the helix by an as-yet undefined allosteric mechanism. Since residues involved in the CusB-CusF interaction[44] have been identified at this end of the barrel it is possible that metal binding stabilises a SilB binding site to aid docking of the metallochaperone to the SilABC efflux complex.

## CONCLUSION

Ag(I) is a potent anti-microbial and a major part of its bactericidal action arises from its ability to mimic Cu(I) as a preferred binding partner to copper binding proteins. Our studies show that key changes in the flexibility of SilF shown by solution based techniques, but not by crystal structure analysis indicates that our crystallisation conditions impact on the helical content of the protein. The differing role of water in the two binding mechanisms allows an adaptability of the binding site to accommodate the two different cations with similar affinity.

The efficiency with which Ag(I) can be removed from bacteria is essential for biological function in these organisms as this ion serves no biological purpose. Evidence is now emerging that carriage of *sil* genes, either on the chromosome or via a plasmid, is not a pre-requisite for Ag(I) resistance. It is a mutation in the *silS* gene[15, 45] leading to an upregulation of the *sil* genes that then leads to the observed enhanced resistance. Hence, with our observations of the similar affinity for both cations, it is an increase in expression of the Sil proteins that leads to the observed Ag(I) resistance, rather than simply the expression of proteins that have a higher affinity for Ag(I). Our findings therefore logically lead to the hypothesis that there is not anything particularly novel about the Sil proteins compared with the Cus proteins: resistance simply arises from there being more Sil proteins available to bind Ag(I) in resistant strains compared to non-resistant strains: the *sil* genes are frequently found with the *pco* genes on a Tn7-like mobile genetic element.

Given the results reported here, this leads to an intriguing and more general hypothesis that there is no specific Ag(I) resistance mechanism, there is a Cu(I) resistance mechanisms and metallochaperone function that can accommodate both Ag(I). Our increased release of Ag(I) into the environment[4–7] and the apparent observed rise in Ag(I) resistance[41, 46, 47] has now to be looked at in a different light. This now appears to arise from the lack of discrimination between Cu(I) and Ag(I) by the existing Cu(I) resistance mechanisms, rather than the evolution of a new mechanism. The consequences of this for how we define a particular metal resistance mechanism and understand their evolution in the environment are therefore profound[48].

## ASSOCIATED CONTENT

### Supporting Information

Supporting information:

Materials and Methods, data files and Supporting figures and tables available as a PDF.

## AUTHOR INFORMATION

### Author Contributions

RML performed the cloning, expression, purification and crystallization of SilF, supervised by AQ, SBC and DJS. MH performed the MD and QM/MM simulations. RML and GH performed the AUC and SEC-MALS experiments and RML, GH and DJS analyzed the data. Protein crystallographic data was collected and analyzed by RML, AQ and SBC. ITC was performed and analyzed by JJAGK and RML. EH carried out the HDX-MS experiments and EH and JB analyzed the data. SR-CD measurements were performed by staff on B21 of Diamond Light Source. MD and QM/MM data was analyzed by CMJ, AC and MH. JH and DJS interpreted the data in terms of the biology of the system. The manuscript writing was contributed by all of the authors. All authors have given approval to the final version of the manuscript. / ‡These authors contributed equally.

## ACKNOWLEDGMENT

DJS, SBC, AQ and RML are grateful to the Research Complex at Harwell (UK) for hosting this work and providing access to equipment and facilities. The authors gratefully acknowledge the University of Nottingham High Performance Computing facility for computational resources.

## ABBREVIATIONS

ITC: Isothermal Titration Calorimetry
SEC-MALS: Size Exclusion Chromatography coupled to Multi-Angle Light Scattering
AUC: Analytical Ultracentrifugtion
QM/MM: Quantum Mechanical Molecular Modelling

