## Supplementary Files for "The adaptability of the ion binding site by the Ag(I)/Cu(I) periplasmic chaperone SilF"

^a^School of Biosciences, University of Nottingham, Sutton Bonington Campus, Leicestershire, LE12 5RD, United Kingdom. ^b^Membrane Protein Laboratory, Diamond Light Source, Rutherford Appleton Laboratory, Didcot, Oxfordshire, OX11 0FA, United Kingdom. ^c^Diamond Light Source, Diamond House, Rutherford Appleton Laboratories, Didcot, Oxfordshire, OX11 0FA, United Kingdom. ^d^Research Complex at Harwell, Rutherford Appleton Laboratory, Didcot, Oxfordshire, OX11 0FA, United Kingdom. ^e^Department  of Chemical and Environmental Engineering, University of Nottingham, University Park, Nottingham NG7 2RD. United Kingdom. ^f^Department of Chemistry, University of Oxford, Oxford, Oxfordshire, OX1 3QU, United Kingdom.

1. List of Supplementary Files
2. Materials and Methods
3. Supplementary Tables
4. Supplementary Figures
1. **SUPPLEMENTARY FILES**

**Protein Databank Codes**

- Apo-SilF: 8BBZ
- Ag(I) bound SilF: 8BHU
- Cu(I) bound SilF: 8BWV

**Simulation Files**

All simulation files are freely available online at:

<https://doi.org/10.6084/m9.figshare.21285264.v1>

data consists of:

Computational supporting data consisting of:

- Molecular dynamics and QM/MM input, parameter, coordinates, and parameter files
- Force field parameterisation files
- run-scripts for setup and production runs
- coordinate output files for QM/MM calculations
- analysis scripts and results

2. **MATERIALS AND METHODS**

**Cloning, Expression and Purification**

The SilF gene from *E.coli* (Uniprot: A0A3T0VBZ2) was synthesised by Twist Bioscience (San Francisco, USA) as a gene fragment. Analysis of the construct indicated the first 38 amino acids formed a disordered region, therefore a truncated version (SilF_38-120_) from V38 was PCR amplified from the full-length sequence. The amplified gene was cloned into a pOPINF vector (PPUK, Rosalind Franklin Institute, UK) using PPUK’s in-fusion method, the vector contains an N-terminal His6-tag with a HRV3C cleavage site[1].

pOPINF-SilF_38-120_ was transformed into BL21 (DE3) *E. coli* cells for expression. Overnight precultures were prepared using a single colony grown in 20 mL TB media supplemented with 20 µL of ampicillin (50 mg/mL), grown at 25 °C. Precultures were used to inoculate 1 L of TB media, supplemented by 1 mL of ampicillin (50 mg/mL), using 10 mL of preculture per litre. Cultures were grown at 37 °C with shaking until an OD (600 nm) of 1.2 was achieved, whereupon 1 mL of 1M isopropyl β-D-1-thiogalactopyranoside (IPTG) was used to induce, the cultures were left to grow at 22 °C for 16 h.

Cells were harvested at 5000 × *g* for 10 minutes with the subsequent pellets re-suspend in lysis buffer (50 mM HEPES, 500 mM NaCl, 20 mM imidazole, pH 7.8), using 50 mL per 10 g pellet. Lysis buffer was supplemented with lysozyme (0.1 mg/mL), DNase I (0.1 mg/mL) and 1 Roche cOmplete protease-inhibitor cocktail tablet (EDTA-free). Cells were lysed using a cell disruptor with two passes at 28kpsi, the resulting lysate was centrifuged at 40,000 × *g* for 30mins. The supernatant was run down a 5 mL Ni^2+^ His-Trap column pre-equilibrated with lysis buffer using a peristatic pump, after loading the column was washed with 20 CV wash buffer (50 mM HEPES, pH 7.8, 500 mM NaCl, 50 mM imidazole). A gradient elution was used to elute SilF using Buffer A (50 mM HEPES, 500 mM NaCl pH 7.8) and Buffer B (50 mM HEPES, 500 mM NaCl, 750 mM imidazole pH 7.8), the length of elution was 20 CV with an end concentration of 100% Buffer B, 3 mL fractions were collected.

Fractions containing SilF_38-120_ were pooled together for dialysis, 3C protease and β-mercaptoethanol (5 mM end conc) were added to the sample. Samples were dialysed (Spectra/Por™3 RC Tubing, MWCO 3.5 kDa) overnight against SEC buffer (25 mM HEPES, pH 7.8, 150 mM NaCl). Dialysed material was run down a reverse IMAC Ni^2+^-HisTrap, with samples coming off in the flow through and partially in the wash. SilF was concentrated down to a volume of 1.5ml and loaded onto a HiLoad 16/600 Superdex 75 PG column, pre-equilibrated with SEC buffer, with 3mL fractions collected. SilF_38-120_ presence was confirmed by SDS-PAGE analysis (with approximately 95% purity), with fractions containing the 8.8 kDa protein pooled together and concentrated to 20 mg/mL using a 3 kDa cut-off spin concentrator (Amicon Ultra-15).

**SEC-MALS**

Size exclusion chromatography with multiangle light scattering (SEC-MALS) was carried out using an AKTA Pure25 (GE Healthcare) fitted with DAWN HELEOS-II 18 angle light scattering detector and a Optilab T-rEX refractive index monitor (both Wyatt Technologies). SilF_38-120_ was applied to a Superdex 75 10/300 increase (GE Healthcare) pre-equilibrated with SEC buffer, 100 µL of sample was run at 2 mg/ml. Data was collected and analysed using Astra v7 (Wyatt, California, USA).

**Analytical Ultracentrifugation (AUC)**

AUC sedimentation velocity (AUC-SV) experiments were carried out using the Beckman Optima analytical ultracentrifuge (Beckman, Indianapolis, US). Samples were loaded into 12 mm double sector epoxy resin cells with sapphire windows at both ends, encased in an aluminium cell. Sample volumes were 400 µL reference SEC buffer and 396 µL solute. Concentrations of SilF_38-120_ used were as follows: 2.0 mg/mL, 1.0 mg/mL, 0.5 mg/mL and 0.25 mg/mL. Ag(I) or Cu(I) was added to samples to a concentration of 10mM for holo bound runs. The experiment was performed at 20 °C with a rotor speed of 50,000 rpm for 18 h to ensure complete sedimentation of protein. Radial scans were obtained using absorbance and Rayleigh interference optics with measurements made every 20 s and 120 s respectively.  Data was analysed in SEDFIT[2] using the continuous distribution (c(s)) method. Sedimentation coefficient and molecular mass are determined normalised to buffer density and viscosity at 20 °C. Buffer density and viscosity measurements were made using an Anton Paar DMA5000 with online viscometer.

**Isothermal Titration Calorimetry (ITC)**

ITC experiments were conducted using a MicroCal PEAQ-ITC calorimeter (MalvernPanalytical, UK) in a Coy anaerobic chamber (<5 ppm O_2_). Purified SilF_38-120_ was dialysed into water (for Ag(I) studies) or 1 M NaCl (for Cu(I) & Cu(II)) overnight, then diluted in each respective buffer to 25 µM before being injected into the sample cell. Metal titrants of 250 µM AgNO_3_ and 250 µM Cu(I)/Cu(II) were made in water and 1 M NaCl respectively.

Injections of 1 µL metal titrant were spaced every 2 minutes for a total of 39 injections, with an initial injection of 0.4 µL, stirring of the cell was conducted at 750 rpm at a constant 25 °C. Data analysis was carried out using Microcal PEQA-ITC Software (version 1.40, Malvern Panalytical)..

**Crystallisation and Structure Refinement**

Purified SilF_38-120_ in SEC buffer was screened in several commercially available screening condition kits (SG1 and Morpheus (Molecular Dimensions)), using a protein concentration of 20 mg/mL. Screens of SilF_38-120_ were prepared both without (apo) and with (holo) Ag(I) (in the form of 5 mM AgNO_3_). Crystallisation was carried out using the sitting drop method in CrystalQuick X plates (Grenier, Austria), with 1nL drops mixed with 1nL crystallisation matrix left at 20°C.

Crystals of apo- SilF_38-120_ formed within a couple of days in several conditions with the condition from the SG1 screen condition G3 (0.01 M ZnSO_4_, 0.1 M MES (pH 6.5), 25% v/v PEG 550 MME) opted for use. Further crystal optimisation around this condition was conducted with crystals used from the final condition 0.01 M ZnSO_4_, 0.1 M MES (pH 7.6) 14% PEG 550 MME. Crystals were soaked in cryo-protectant for 30 s then flash frozen in liquid nitrogen and stored.

Crystals of Ag(I)- SilF_38-120_ formed in SG1 screen D10 (0.2 M LiSO_4_, 0.1 M Bis-Tris pH 6.5 and 25% w/v PEG 3350) after approximately a week. Crystals were picked and soaked in cryo-protectant (supplemented with 27% PEG 3350, 15% glycerol & 5 mM AgNO_3_) for 30 s before flash frozen in liquid nitrogen and stored. Crystals of Cu(I)- SilF_38-120_ also formed in the SG1 screen however this time condition C4 (0.2 M potassium sodium tartrate tetrahydrate & 20% w/v PEG 3350). Crystals grew after approximately 2 weeks in anaerobic conditions, the crystals were picked and soaked in cryo-protectant supplements with 27% PEG 3350, 15% glycerol and 5 mM CuCl.

X-Ray diffraction data was collected on beamline I24 at Diamond Light Source (Oxfordshire, UK). The structures were solved by molecular replacement in Phaser[3]. The apo structure was solved using CusF (PDB 2BV3) as a model, all subsequent structures were solved using the apo-SilF_38-120_ structure as the model. Further model building and refinement were carried out in Coot[4] and Refmac[5] (version 5.8.0258) respectively, refined models were evaluated through MolProbity[6]. X-Ray and refinement data is given in Table S2.

**Hydrogen-Deuterium Exchange Mass Spectrometry (HDX)**

For initial peptide mapping, SilF_38-120_ (at 40 µM) was diluted ×11 in buffer E (20 mM HEPES, 30 mM KNO_3_, pH 7.8) and quenched 1:1 with 100 mM KH_2_PO_4_/K_2_HPO_4_, 2 M GuHCL, pH 2.08. 50 µL of this sample was injected into a Waters HDX Manager with an immobilized pepsin column (2.1 × 30 mm; Waters), C18 trapping column (VanGuard ACQUITY BEH 2.1 × 5 mm; Waters), and analytical C18 column (1.0 × 100 mm ACUITY BEH; Waters). Mass spectrometer – Synapt G2-Si. Mobile phases were 0.1% formic acid in H_2_O (A) and 0.1% formic acid in ACN (B), such that their pH was 2.55. Protein was applied to the pepsin and trapping columns in A at 100 μL/min and eluted from the analytical column according to the following elution profile  using H_2_O/ACN (+0.1% formic acid v/v): 1 – 7 minutes 97% water to 65% water, 7 – 8 minutes 65% water to 5% water, 8 – 10 minutes held at 5% water.

Sample preparation of SilF_38-120_ in its apo and Ag(I) bound states for labelling experiments were conducted in the same manner as mapping, however sample buffer E was made in D_2_O instead of H_2_O (buffer L) and quenching occurred after 30 s, 5 minutes and 30 minutes.

Peptide sequences were assigned from MSE fragment data with Protein Lynx Global Server 3.0.3 (Waters) and DynamX 3.0 (Waters). Labelling data was acquired as for sequencing, except the mass spectrometer acquired MS scans only. Differences in uptake were filtered using hybrid significance testing using Deuteros 2.0 and overlaid on the SilF_38-120_ protein structure using Pymol.

**Synchrotron Radiation Circular Dichroism (SR-CD)**

CD experiments were performed on beamline B23 of the Diamond Light Source using a nitrogen-flushed ChirascanPlus CD spectropolarimeter (Applied Photophysics Ltd, Leatherhead, UK). Five samples, all of 5 mg/mL concentration (and their corresponding buffers) were supplied: one native peptide, SilF apo, (buffer: HEPES 20 mM, KNO_3_ 30 mM), one peptide at pH 5 (buffer: CH_3_COONa 20 mM, KNO_3_ 30 mM), one at pH 9 (buffer: bicine 20 mM, KNO_3_ 30 mM), one peptide with Cu(I) ions (1:10 ratio, buffer: HEPES 20 mM, KNO_3_ 30 mM, 10 Cu(I) equivalent) and one peptide with Ag(I) ions (1:1 ratio, buffer: HEPES 20 mM, KNO_3_ 30 mM, 1 Ag(I) equivalent).

The samples were studied across two regions: near-UV (250-330 nm) and far-UV (180-260 nm). The measurements were acquired using an integration time of 1 s, cuvettes of 0.002 cm (demountable – for far-UV) and 0.2 cm pathlength cuvette (for near-UV) with 1 nm bandwidth at 25 °C. Four repetitions were acquired for each sample. The data obtained was processed using CDApps – for the far-UV region[7] and OriginLab.

**Computational methods**

**Parametrization.** Since the recent non-bonded set of classical parameters failed to correctly describe the metal coordination in the molecular dynamics (MD) simulation of SilF_38-120_ protein, we generated bonded force field parameters for Cu(I) and Ag(I) ions using Seminario/ChgModB method available through the Python module of Metal Centre Parameter Builder (MCPB.py) in Amber18 software. We used X-Ray structures of Cu(I)- SilF_38-120_ and Ag(I)- SilF_38-120_ which both have one histidine and two methionine residues in their metal coordination sites. We performed the geometry optimization and force constant calculations for the sidechain model and the Merz-Kollman RESP charge calculation for the large model using B3LYP/def2-TZVP level of theory in Gaussian16 program. The Lenhard-Jones parameters for monovalent cations Cu(I) and Ag(I) were obtained from[8]. Final force field parameters are available in the Supporting Information (SI) detailed above. We used the H++ webserver to determine the protonation states of the titratable residues in SilF_38-120_ protein using the physiological conditions (pH = 7, Salinity = 0.15, Internal Dielectric = 10, External Dielectric = 80). The ff14SB force field parameters were used to model the standard protein residues. The final systems were solvated in the truncated octahedron of TIP3P water molecules (10 Å from the solute) and neutralized adding chloride counterions. The Apo form of SilF_38-120_ protein without a metal present was modelled at the physiological (pH = 7) conditions using a similar approach.

**Classical MD simulations.** Following the initial 1000 steps of solute-restrained (20 kcal mol^-1^) steepest descent minimization we further relaxed the systems by performing Langevin MD simulations at the constant temperature (300 K with a collision frequency of 2 ps-1) and pressure (1 atm with a relaxation time of 2 ps using the Berendsen barostat) applying the equivalent weak positional harmonic restraints on protein atoms for a total of 1 ns. After a short relaxation, the systems were subject to four parallel production *NPT*simulations (1.6 ms each) using 2 fs time step and SHAKE algorithm recording a snapshot every 2 ps. Periodic boundary conditions were applied in all directions while Particle Mesh Ewald method with a cutoff of 12 Å was used to account for long-range electrostatics. All MD simulations were carried out using *pmemd* module while the analysis of the resulting trajectories was performed using *cpptraj* tools of Amber18 software.

**QM/MM simulations.** A set of previously obtained snapshots were further minimized for 300 and 200 steps of steepest descent and conjugated gradient unrestrained minimization respectively, using a coupled QM/MM potential. While a classical ff14SB force field has been chosen to describe MM atoms, the B3LYP/def2-TZVP method was used to treat QM atoms. The metal ion, the sidechains of two Met and one His residues were treated quantum mechanically (QM). Minimized structures were subject to 2 ps of *NVT* equilibration before collecting the additional production simulation data for total of 40 ps recording a snapshot every 2 fs. A similar simulation setup has been employed as described earlier. All water molecules, counterions and the rest of the protein was modelled classically. All QM/MM calculations were carried out using *sander* module of Amber18 program with the QM calculations performed externally employing Gaussian 16 package.

**ONIOM calculations.** To calculate the binding affinity, we extracted the suitable snapshots from QM/MM simulations and performed the two-layer ONIOM calculations using a full SilF protein in the presence and the absence of the metal (see **Figure 2**). The residues found within 5 Å from the QM zone were allowed to move freely during the optimization, and nearest 200 water molecules around the QM region were retained. The metal and the sidechains of the coordinating residues were described with QM while the rest of the protein, solvent and counterions was treated using classical MM. The optimizations and frequency analysis were carried out using the ONIOM[B3LYP-D3/def2- TZVP:ff14SB] level of theory employing the mechanical embedding followed by the electronic embedding single point calculation at the same level of theory. All ONIOM calculations were carried out with Gaussian16 software. The solvation enthalpy of metal ions was obtained using the implicit SMD single point calculations.

**SUPPLEMENTARY TABLES**

**Supplementary Table 1: SEC-MALS analysis of SilF_38-120_**

| SEC-MALS Output | Values |
| --- | --- |
| Radius of hydration (rh(Q)z) (nm) | 2.323 (±8.365%) |
| Average rh(Q) (nm) | 1.884 (±1.670%) |
| Number averaged molecular weight (Mn) (g/mol) | 8.743x10^3^ (±7.888%) |
| Mp (g/mol) | 8.664x10^3^ (±7.082%) |
| Weight averaged molecular weight (Mw) (g/mol) | 8.769x10^3^ (±8.116%) |
| Polydispersity (Mw/Mn) | 1.003 (±4.398%) |

**Supplementary Table 2: Protein crystallography refinement statistics**

|  | apo-SilF | Ag(I)-SilF | Cu(I)-SilF |
| --- | --- | --- | --- |
| Wavelength | 0.999 Å | 0.999 Å | 0.999 Å |
| Resolution range | 45.95 - 2.2 (2.279 - 2.2) | 51.41 - 1.7 (1.761 - 1.7) | 46.92 - 2.2 (2.279 - 2.2) |
| Space group | P 65 2 2 | P 21 21 21 | I 21 21 21 |
| Unit cell | 109.47 109.47 84.59,  90 90 120 | 60.93 81.59 95.78,  90 90 90 | 77.29 77.29 187.69,  90 90 90 |
| Total reflections | 572835 (46562) | 713161 (65325) | 701243 (47107) |
| Unique reflections | 15524 (1502) | 53221 (5226) | 28977 (2855) |
| Multiplicity | 36.9 (31.0) | 13.4 (12.5) | 24.2 (16.5) |
| Completeness (%) | 98.87 (97.85) | 99.95 (100.00) | 99.90 (99.40) |
| Mean I/sigma(I) | 15.1 (1.0) | 12.8 (0.7) | 7.8 (1.8) |
| Wilson B-factor | 46.35 | 26.62 | 43.13 |
| CC1/2 | 1.0 (0.6) | 0.998 (0.328) | 0.997 (0.901) |
| Reflections used in refinement | 15521 (1502) | 53213 (5226) | 28904 (2852) |
| Reflections used for R-free | 802 (74) | 2618 (274) | 1420 (150) |
| R-work | 0.2154 (0.2819) | 0.2025 (0.2823) | 0.2794 (0.3935) |
| R-free | 0.2493 (0.3449) | 0.2335 (0.2584) | 0.3197 (0.4211) |
| Number of non-hydrogen atoms | 1805 | 3790 | 3655 |
| macromolecules | 1792 | 3563 | 3634 |
| ligands | 2 | 60 | 12 |
| solvent | 11 | 167 | 9 |
| Protein residues | 235 | 469 | 478 |
| RMS(bonds) | 0.014 | 0.015 | 0.010 |
| RMS(angles) | 1.94 | 1.84 | 1.21 |
| Ramachandran favored (%) | 98.69 | 99.78 | 94.21 |
| Ramachandran allowed (%) | 1.31 | 0.22 | 5.58 |
| Ramachandran outliers (%) | 0.00 | 0.00 | 0.21 |
| Rotamer outliers (%) | 2.03 | 0.00 | 1.75 |
| Molprobity score | 1.50 | 1.21 | 2.40 |
| Average B-factor | 53.13 | 33.59 | 53.66 |

**Supplementary Table S3: Bond lengths for Ag(I) and Cu(I) bound to SilF and CusF, respectively.**

| **Bond**  **(X = metal ion)** | **SilF-Ag(I) (Å)** | **CusF- Ag(I) (Å)** | **SilF- Cu(I) (Å)** | **CusF- Cu(I) (Å)** | **SilF-Ag(I)**  **QM/MM^1^** | **SilF-Cu(I)**  **QM/MM^1^** | **SilF-Ag(I)**  **Classical** | **SilF-Cu(I)**  **Classical** |
| --- | --- | --- | --- | --- | --- | --- | --- | --- |
| N-X (His60) | 2.3 | 2.2 | 2.3 | 2.0 | 2.3 | 2.1 | 2.3 | 2.0 |
| S-X (Met73) | 2.7 | 2.7 | 2.2 | 2.2 | 2.8 | 2.4 | 2.6 | 2.3 |
| S-X (Met75) | 2.5 | 2.4 | 2.4 | 2.2 | 2.6 | 2.4 | 2.6 | 2.3 |
| CE3/CZ3-X (Trp70) | 2.8/2.9 | 2.8/3.0 | - | 2.7/2.9 | 3.7/4.0 | 4.0/4.3 | 3.3/3.7 | 3.8/4.2 |
| H_2_O-X | - | - | 2.5 |  | 6.6 | 1.8 | 2.7 | 2.1 |

**SUPPLEMENTARY FIGURES**

**Supplementary Figure 1A**

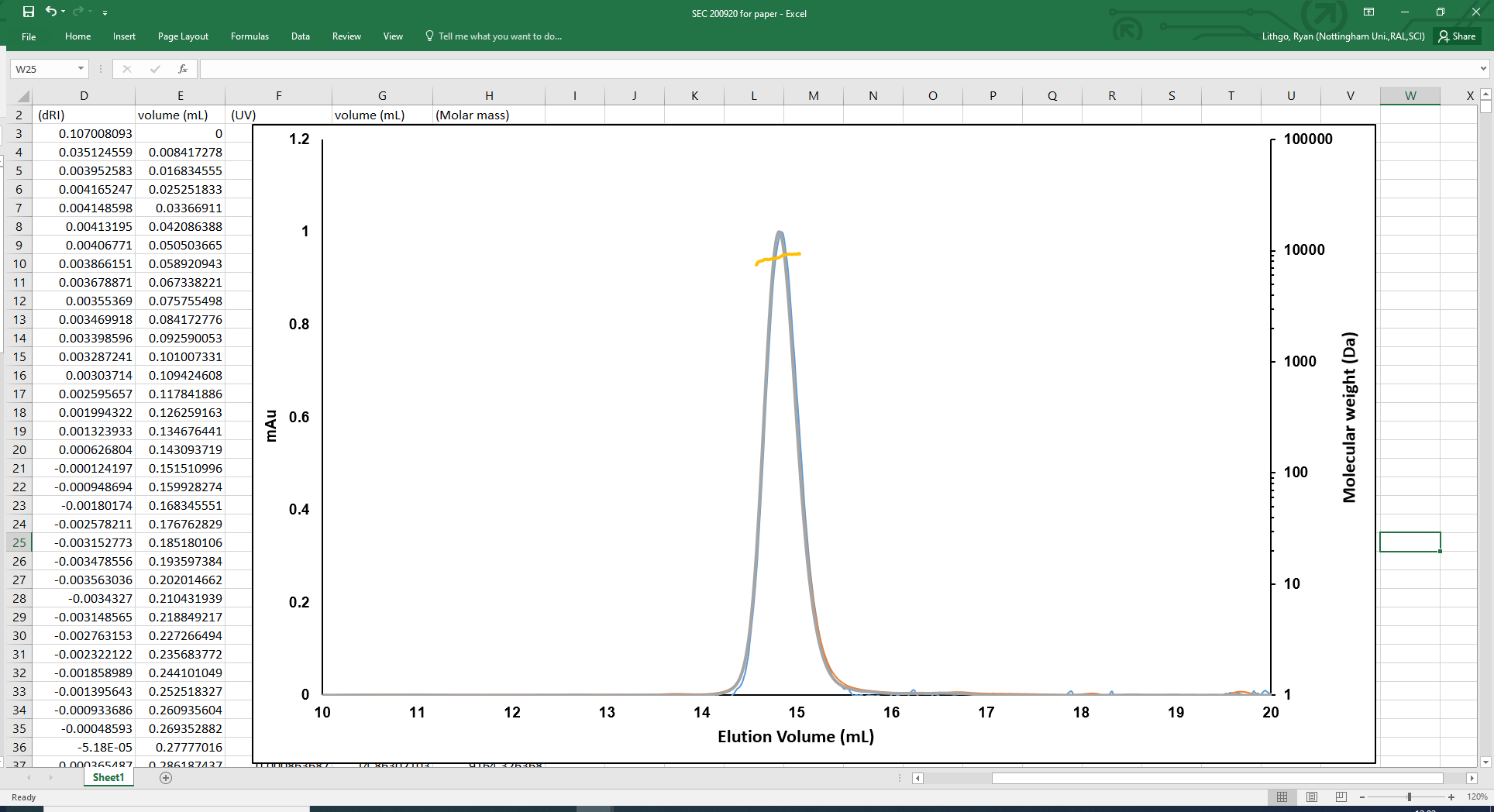

**Supplementary Figure 1B**

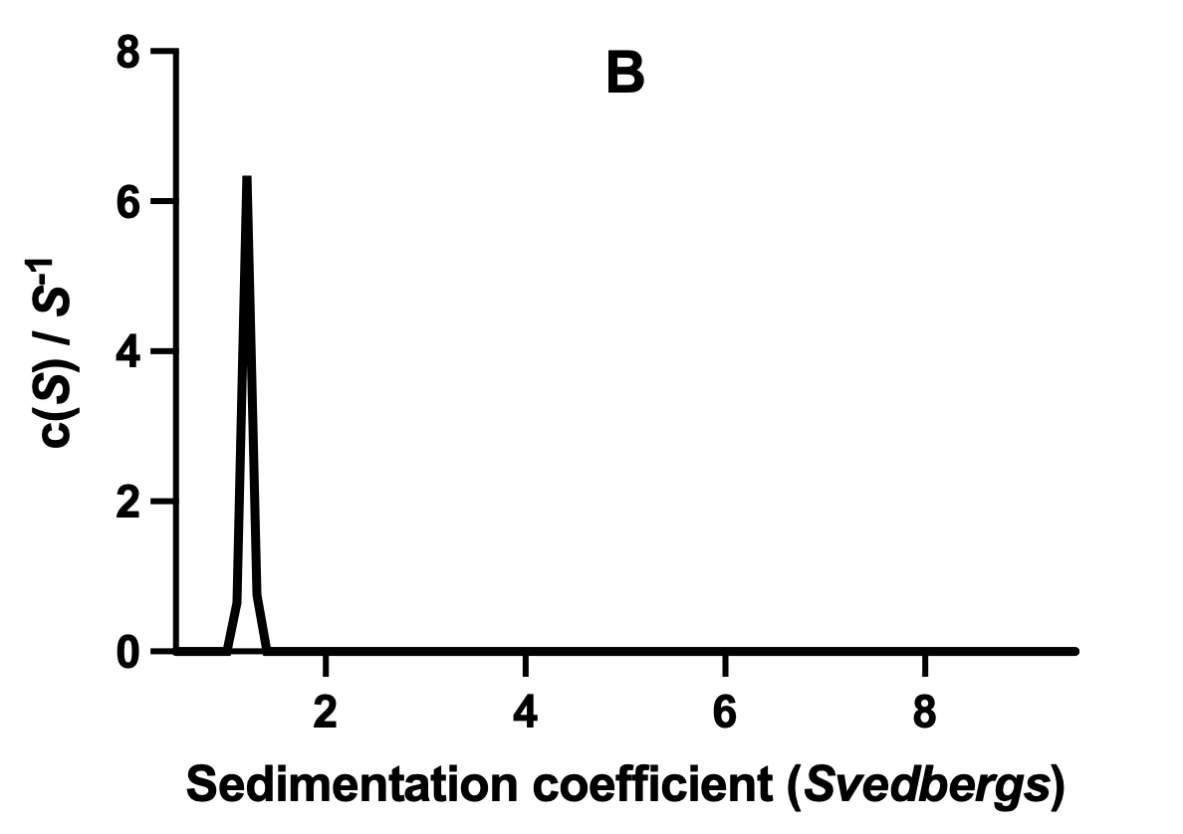

**Figure S1A:** SEC-MALS analysis of SilF. The chaperone elutes as a single peak as judged by the UV absorbance trace taken at 280 nm. The calculated molecular weight across the peak is shown in yellow and corresponds to that expected for a monomer of 9 kDa. **B:** Sedimentation coefficient distribution derived from sedimentation velocity analytical ultracentrifugation showing a single species at 1.25 S with no evidence of higher order aggregation states

**Supplementary Figure 2:**

**­**
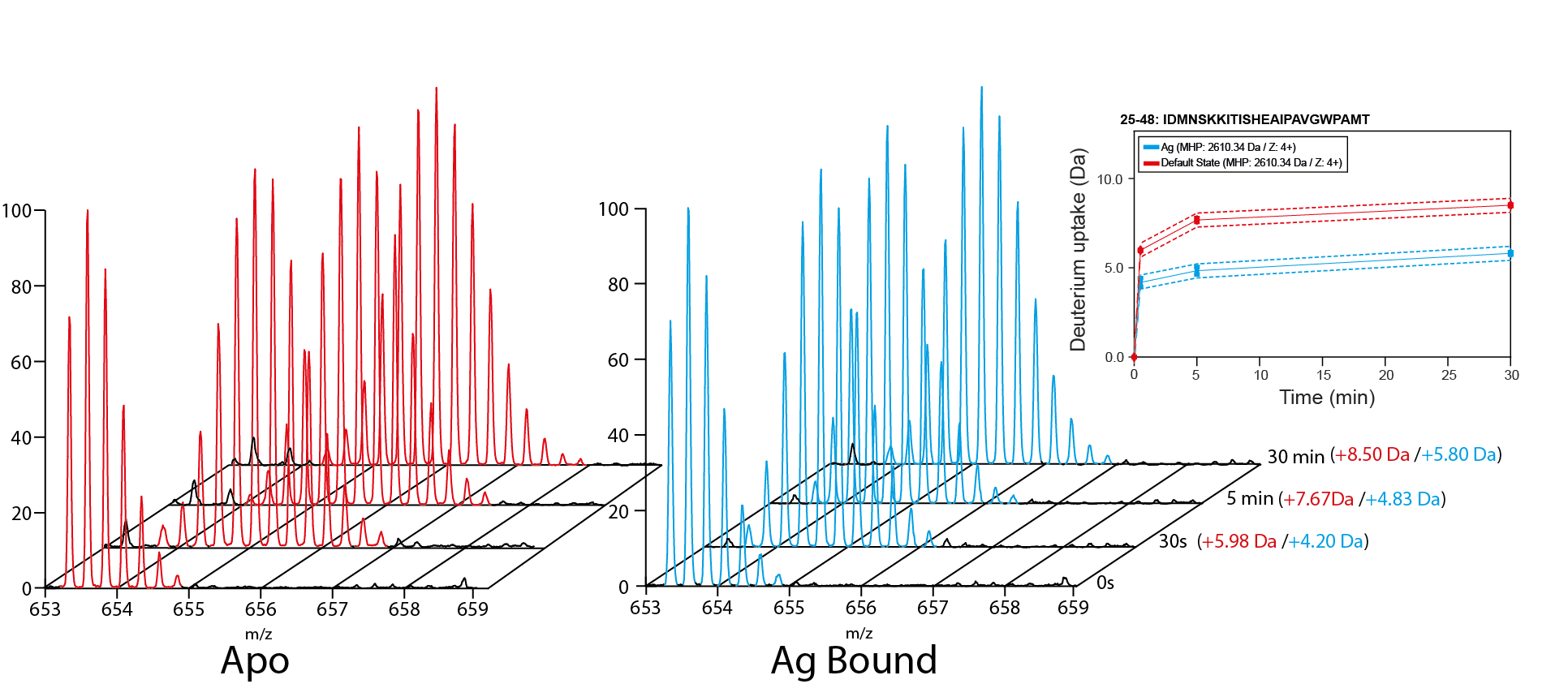

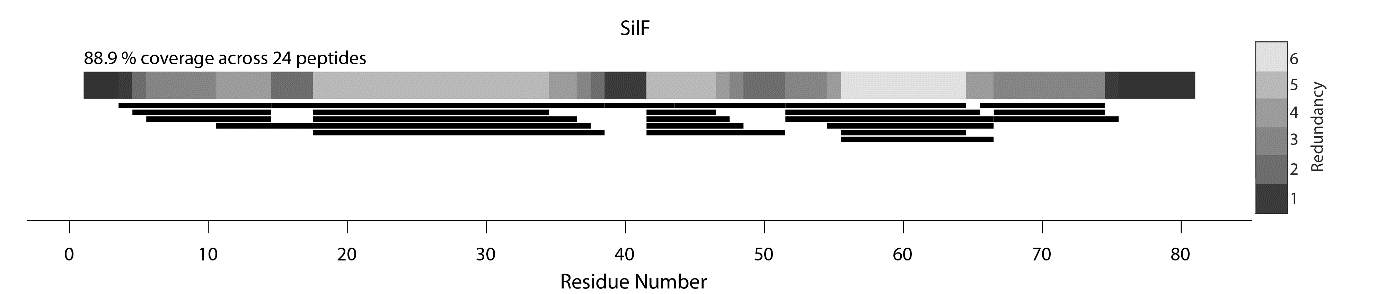

**Figure S2:** Top panels, shows the raw spectra of peptide IDMNSKKITISHEAIPAVGWPAMT (aa25-48) and a combined deuterium uptake plot over time. Apo results in a far greater m/z shift over time than Ag(I) bound, due to the protection of the amino acids interacting with Ag(I) when in the bound state. Bottom panel is a coverage map of the peptic peptides identified along the amino acid sequence of SilF.

**Supplementary Figure 3**

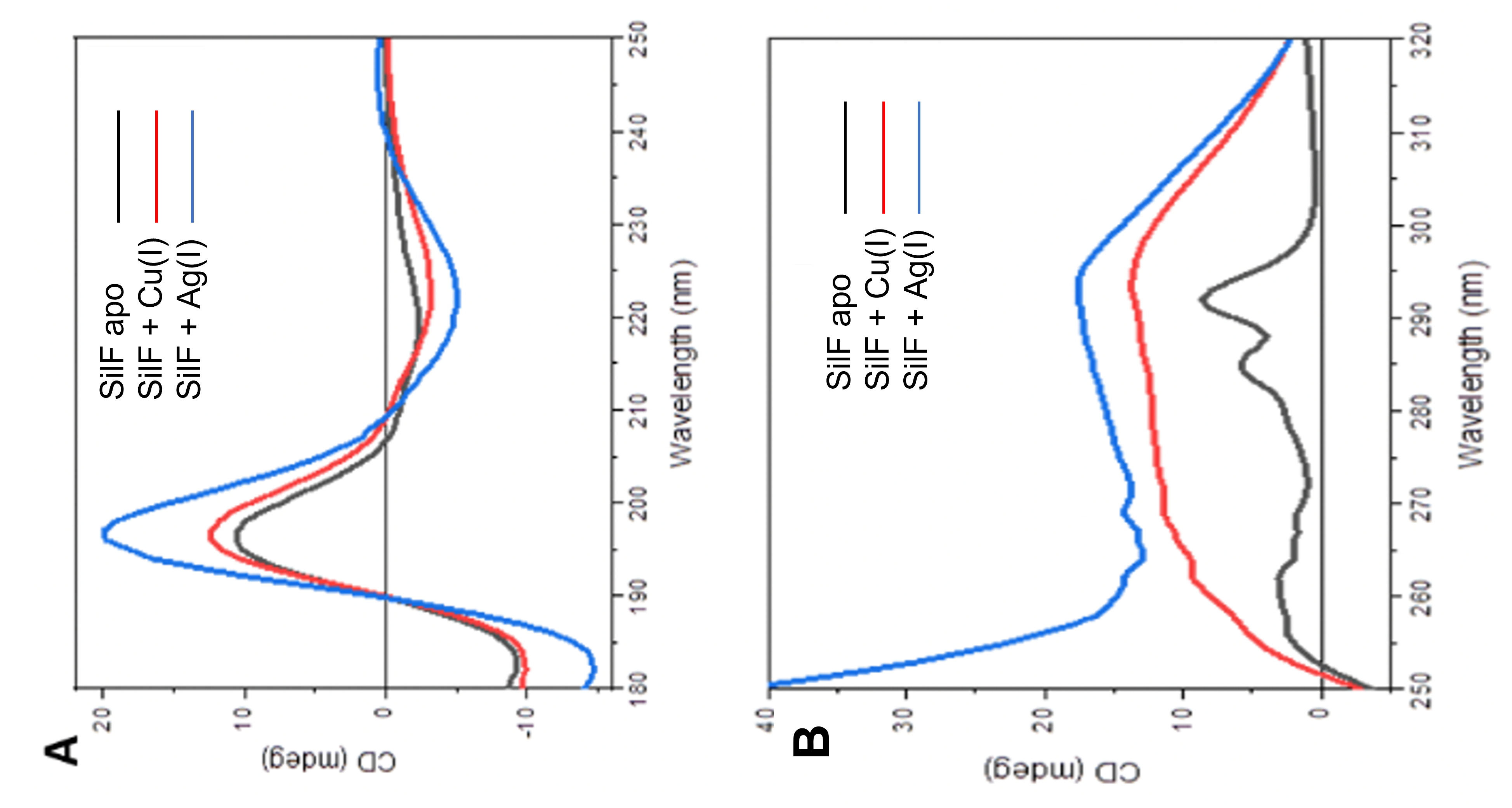

**Figure 4:** Synchrotron Radiation circular dichroism (SRCD) of SilF in the apo, Cu(I) and Ag(I) bound forms. **A:** Far UV SRCD showing changes to secondary structure **B:** Near UV SRCD showing changes to aromatics residues upon ion binding

**Supplementary Figure 4**

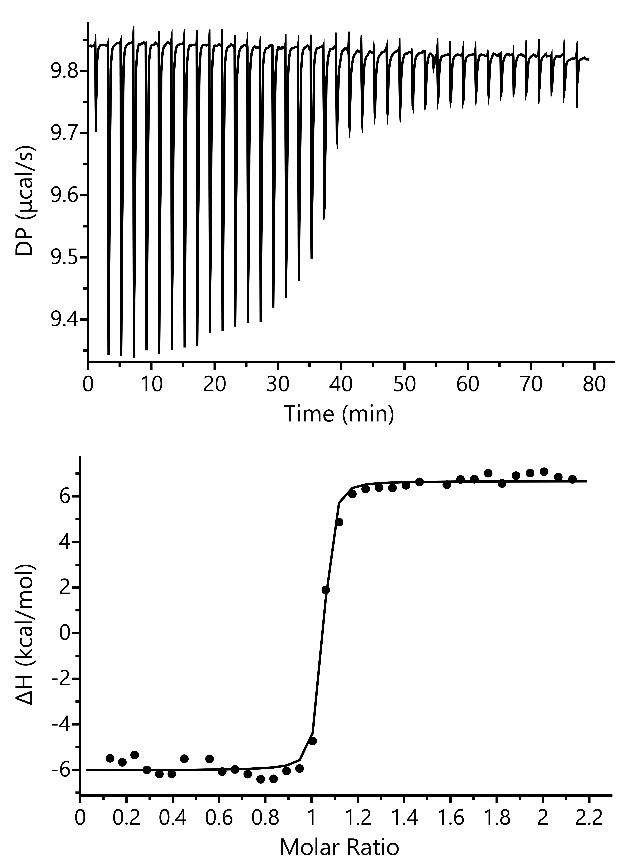

**A**

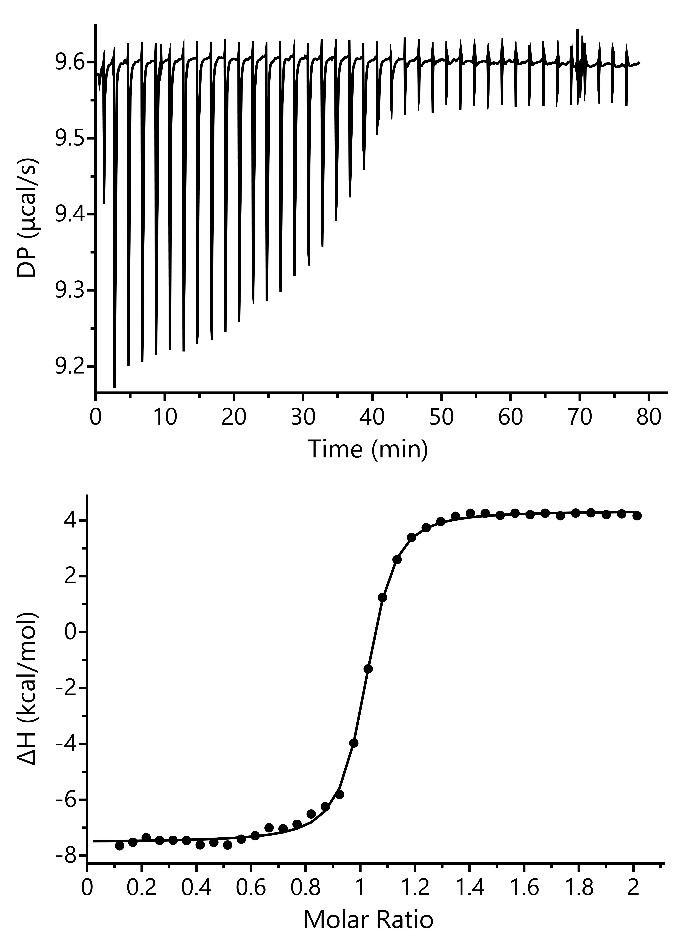

**B**

**Figure S4:** Isothermal titration data for A) Ag(I)-SilF and (B) Cu(I) SilF. The upper panels are the raw thermograms, and the lower panels show data fitted to a 1:1 binding model to the integrated heat data per injection.

**Supplementary Figure 5**

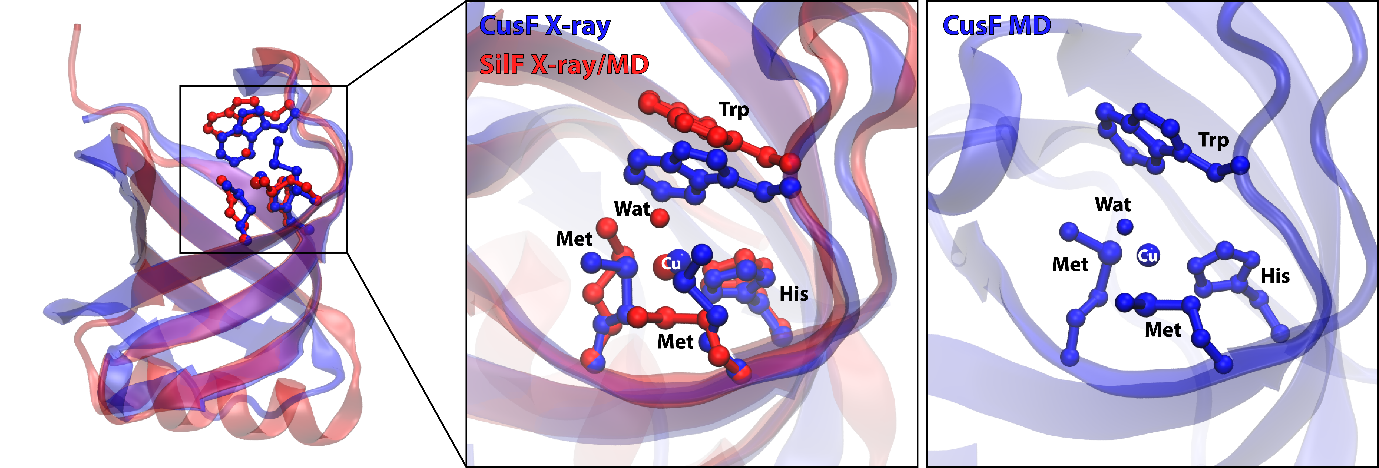

**Figure S5:** Comparison of different binding modes obtained for Cu(I) in CusF and SilF. In X-ray structure of CusF with Cu(I) (blue) the sidechain of Trp is found closer to the copper while the structure of SilF demonstrates the coordination of a bound copper by one water molecule (red). A similar structure is obtained in MD simulations for both CusF and SilF whereby water molecules have strong preference towards coordinating copper in its bound state.
